# The Role of Linker Length and Composition in Actin Binding and Bundling by Palladin

**DOI:** 10.1101/2025.11.06.687016

**Authors:** Rachel A. Sargent, Colby W. Bradford, Lauren M. Hughes, Nathan H. Ta, MarcArthur J. Limpiado, Ravi Vattepu, Joseph G. Brungardt, Moriah R. Beck

## Abstract

Interdomain linkers in multidomain proteins influence spatial arrangement, flexibility, and cooperative binding, yet their functional roles remain underexplored. Palladin, an actin scaffold protein essential for cytoskeletal organization, contains tandem immunoglobulin domains (Ig3-4) separated by an unusually long (∼41-residue), arginine-rich, intrinsically disordered linker. While Ig3 mediates direct F-actin binding, the adjacent Ig4 domain and linker enhance binding and bundling, suggesting active roles beyond passive connection. Using targeted mutagenesis, actin co-sedimentation, bundling assays, and SAXS, we show that linker length, charge, and residue patterning are critical for activity. Shortening or replacing the 41-residue linker with the native Ig4–Ig5 linker reduced binding and abolished bundling, while small internal deletions decreased affinity but unexpectedly increased bundling efficiency. Neutralizing basic residues eliminated activity, increasing net positive charge favored bundling at the cost of binding affinity, and scrambling the sequence impaired both functions—demonstrating that electrostatic tuning and residue order, not just net charge, are essential. SAXS revealed that the native linker supports a broad, dynamic interdomain ensemble, enabling multiple binding-competent geometries. Shortened linkers restricted this conformational space, locking domains in extended orientations incompatible with efficient actin engagement. These results establish the Ig3–Ig4 linker as a finely tuned structural and electrostatic module that coordinates domain flexibility and filament crosslinking. More broadly, these findings highlight long, intrinsically disordered linkers as active determinants of cytoskeletal scaffold function, providing a general mechanism for modulating multivalent interactions in actin-binding proteins.

## Introduction

Immunoglobulin (Ig) domains are among the most abundant and evolutionarily conserved structural motifs in the human proteome, found in diverse proteins ranging from cell adhesion molecules to cytoskeletal scaffolds (1). While these domains share a common β-sandwich fold, their functional versatility is largely driven by how they are arranged and connected in multidomain proteins (2, 3)(4). In tandem Ig domain arrangements, short interdomain linkers (typically 3–6 residues in length) play critical roles in determining the orientation, spacing, and dynamic interplay between adjacent domains. Several prior studies have recognized that these linkers are not passive connectors; rather, they actively influence domain flexibility, folding cooperativity, mechanical response, and interactions with binding partners (5-7). Despite this growing recognition, the molecular rules governing how linker length, sequence, and post-translational modifications (PTMs) shape protein function remain incompletely understood.

Cytoskeletal scaffold proteins offer an excellent context for studying the functional roles of interdomain linkers, as many rely on precise spatial organization and dynamic assembly of modular domains to interact with filamentous actin and associated proteins. Palladin is one such scaffold protein, playing a central role in organizing actin structures during cell migration, adhesion, and morphogenesis. It is widely expressed and upregulated in pathological settings such as cancer cell invasion (8, 9), bacterial infection (10), and neurodevelopmental disorders (11, 12). Palladin’s actin-binding activity is primarily mediated by its tandem Ig3-4 domains, which are separated by an unusually long, ∼40-residue, arginine-rich linker. This linker stands out compared to typical Ig domain linkers in both length and composition and is predicted to be intrinsically disordered (Figure 1). Moreover, it undergoes phosphorylation by Akt1, suggesting a potential regulatory function (13).

**Figure 1.**
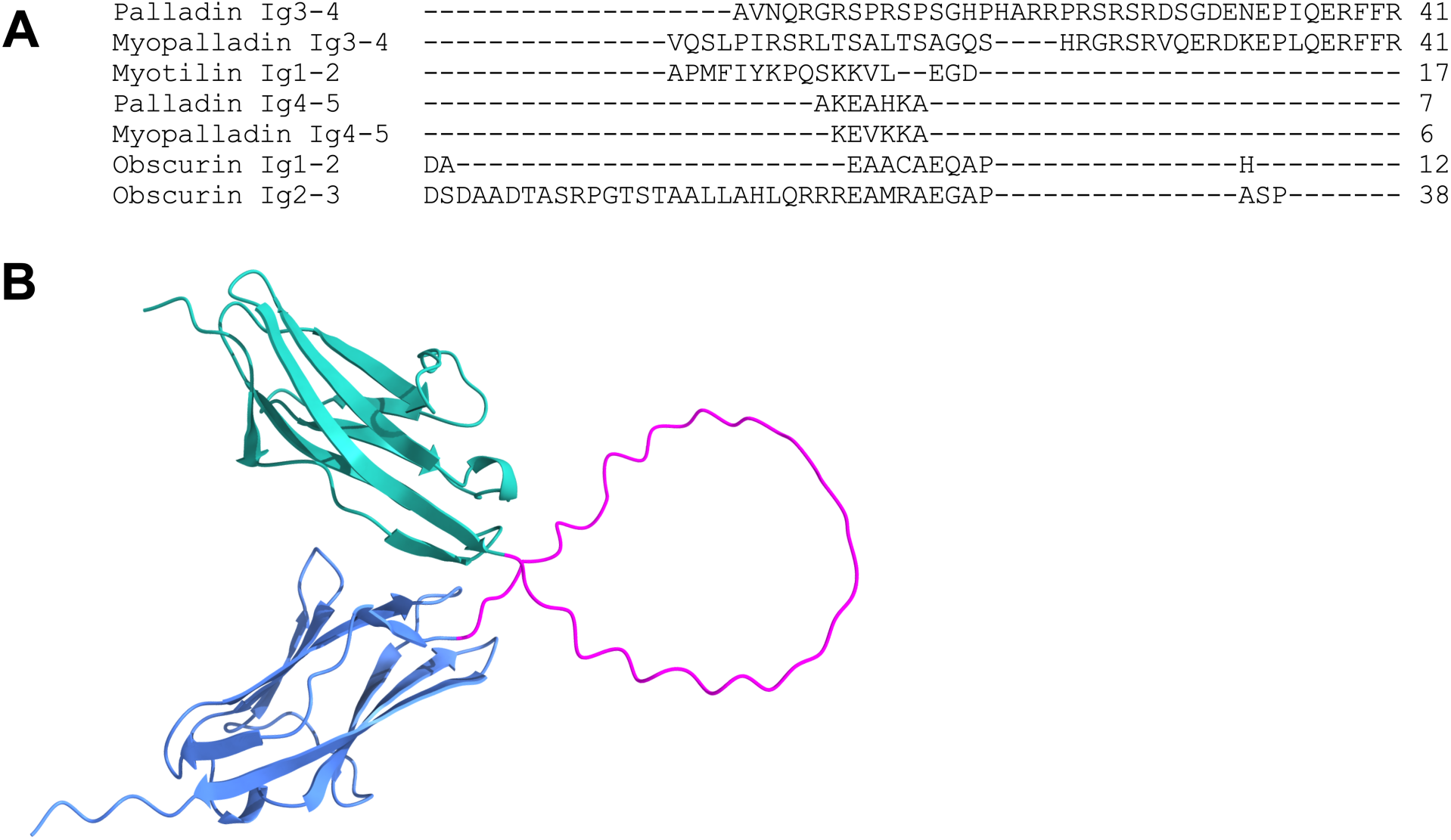
Palladin Ig3-4 Linker sequence alignment and predicted structure. **(A)** Sequence alignment of the interdomain linker connecting tandem Ig domains in palladin (Ig3-4, Ig4-5) compared to corresponding linker regions in myopalladin (Ig3-4 and Ig4-5), myotilin (Ig1-2) and obscurin (Ig1-2 and Ig2-3). The palladin and myopalladin Ig3-4 linkers are notably longer (∼40 residues) and enriched in positively charged residues (arginine and lysine), suggesting potential for electrostatic interactions with F-actin. **(B)** AlphaFold-predicted structure of the palladin Ig3-4 tandem domain with the full 41-residue native linker. While Ig3 and Ig4 are modeled as well-folded Ig-like β-sandwich domains, the interdomain linker is predicted to be intrinsically disordered, lacking defined secondary structure. This unstructured conformation likely imparts flexibility, allowing dynamic interdomain positioning that facilitates multivalent actin binding and bundling.

Previous studies have shown that the Ig3 domain is sufficient for direct F-actin binding and bundling (14), yet the presence of the adjacent Ig4 domain and the extended linker enhance these activities. This raises important questions about how the structural and dynamic properties of the linker contribute to actin remodeling. The unusually long and positively charged linker may facilitate conformational flexibility or serve as a regulatory interface, but its precise functional role has not been defined.

In this study, we use palladin’s Ig3-4 tandem as a model system to investigate the functional consequences of linker variation. Through mutagenesis and structural analysis, we probe how changes to linker sequence and length affect actin binding, bundling, and domain organization. Our findings support the view that linker regions are active determinants of protein function and suggest broader implications for understanding how disordered and modular regions cooperate to regulate cytoskeletal architecture.

## Results

### Linker region influences actin binding and bundling

Previous studies have demonstrated that the tandem Ig3-4 domains of palladin exhibit stronger binding affinity for F-actin than the isolated Ig3 domain (14). However, the ability of the Ig3 domain alone to bundle F-actin has been inconsistently reported. Dixon *et al*. initially concluded that Ig3 was incapable of crosslinking F-actin, whereas Gurung *et al.* later demonstrated that Ig3 can mediate F-actin bundling, particularly at higher concentrations and more effectively under G-actin buffer conditions (15). Notably, neither study examined the role of the linker region connecting Ig3 to Ig4 in modulating actin-binding or bundling activity. To address this gap, we compared the actin-binding affinity and bundling efficiency of multiple palladin constructs: the isolated actin-binding domain (Ig3), the Ig3 domain with its adjacent linker region (Ig3L), the tandem Ig3-4 domains (Ig3-4), and combinations of Ig3 or Ig3L added in trans to Ig4. Actin co-sedimentation assays revealed that Ig3L binds more strongly to F-actin than Ig3 alone, although not as tightly as the covalently linked Ig3-4 construct, indicating that the linker enhances actin binding when physically connected to Ig3 (Figure 2A, Table 1). Bundling assays showed a similar trend: Ig3L increased bundling efficiency compared to Ig3, but remained less effective than Ig3-4 (Figure 2B). Importantly, when Ig4 was added in trans to either Ig3 or Ig3L, no significant enhancement in binding or bundling was observed compared to the individual constructs, suggesting that non-covalent association is insufficient to replicate the cooperative effects seen in the tandem domain. These results collectively demonstrate that bot the linker region and Ig4 domain contribute to palladin’s ability to bind and bundle F-actin, but only when integrated into a single protein chain. The covalent linkage appears necessary to maintain a favorable spatial arrangement and dynamic interplay between domains, enabling multivalent interactions with actin filaments.

**Figure 2:**
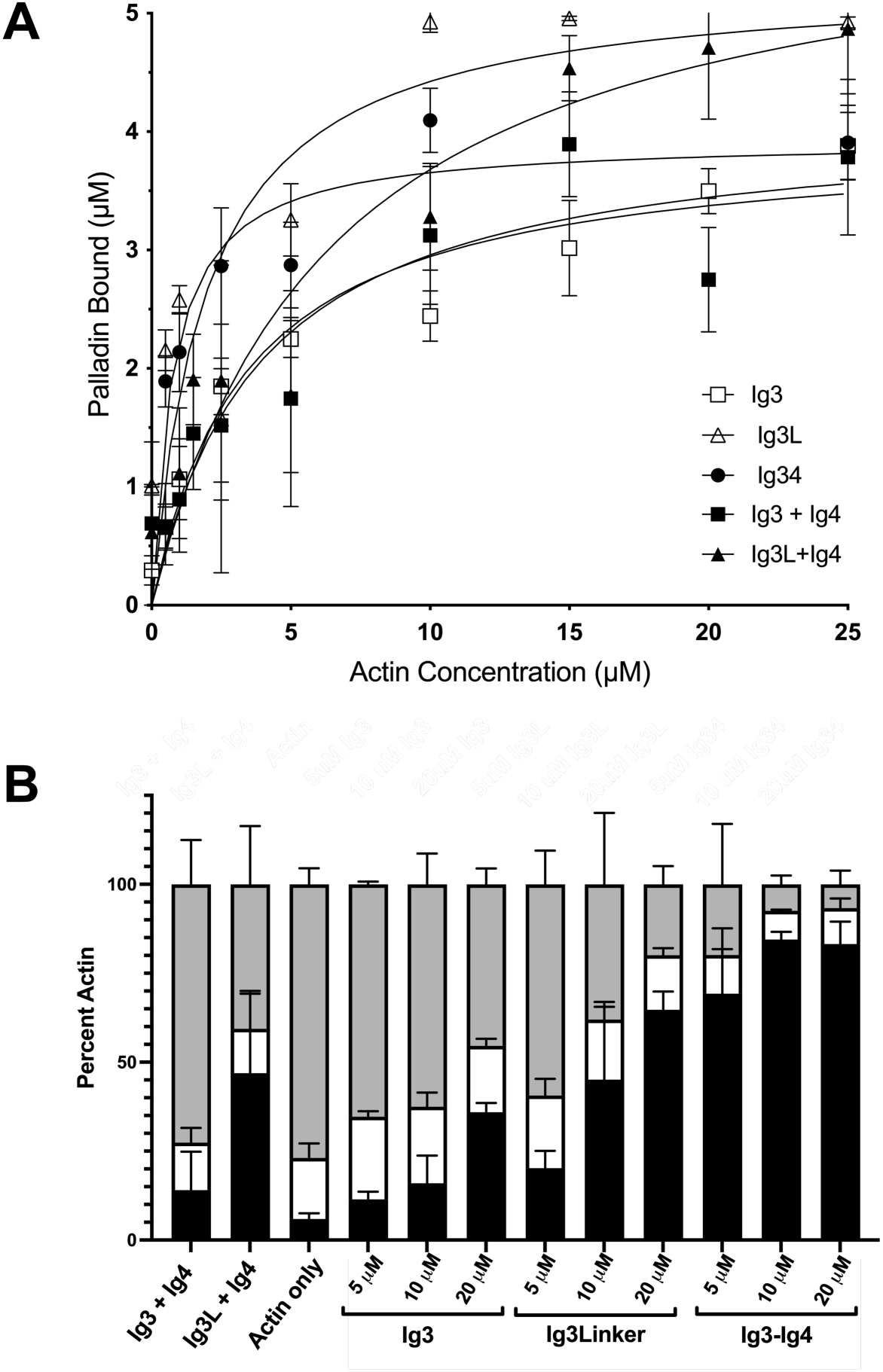
The linker region enhances F-actin binding and bundling activity of palladin Ig domains. **(A)** F-actin binding curves for palladin constructs: Ig3 (open squares), Ig3 with the native linker region (Ig3L, open triangles), tandem Ig3–Ig4 domains (Ig3-4, circles), Ig3 + Ig4 (closed squares) and Ig3L + Ig4 (closed triangles). Binding data were fitted to a hyperbolic binding equation to determine K_d_ and B_max_ values (see Table 1). Inclusion of the linker increases binding affinity compared to Ig3 alone, though not to the level observed whenIg3–Ig4 are covalently connected in a tandem construct. When the domains were added in trans, no enhancement in binding was observed. **(B)** F-actin bundling activity as assessed by low-speed co-sedimentation assay. Samples include: Ig3 + Ig4 (10 μM each), Ig3L + Ig4 (10 μM each), and actin alone (control), followed by individual constructs at increasing concentrations (5, 10, and 20 μM). Bar graphs show the distribution of actin between the pellet (gray), supernatant (white), and bundled F-actin (black portion of the bar). Bundling efficiency increases with protein concentration and is highest for the Ig3–Ig4 tandem construct. The linker region enhances bundling when fused to Ig3, whereas separate addition of Ig4 does not replicate this effect. For all data are means +/- standard deviation for at least three separate measurements.

**Table 1:**
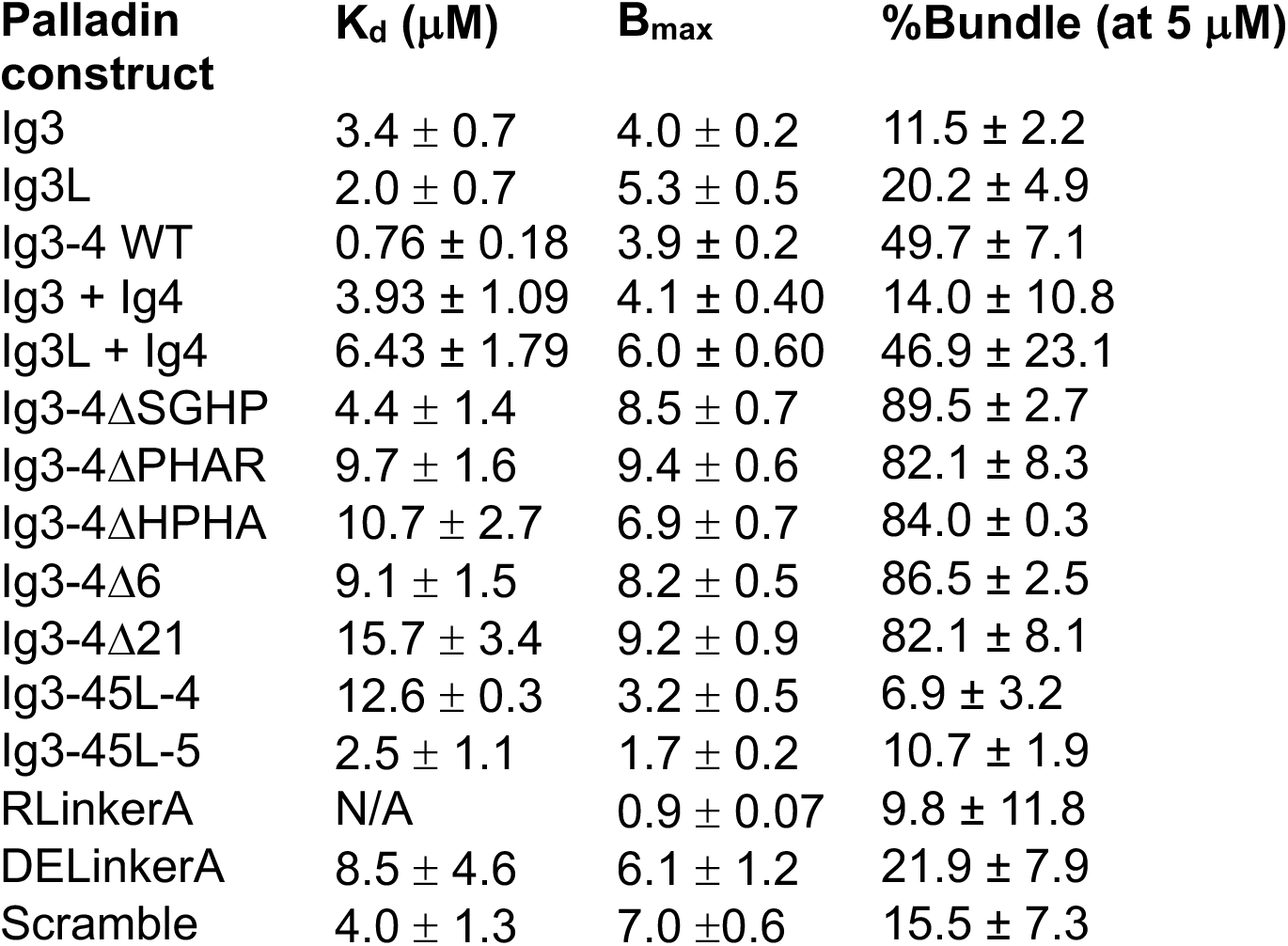
Actin binding data.

| **Palladin construct** | **K<sub>d</sub> (μM)** | **B<sub>max</sub>** | **%Bundle (at 5 μM)** |
| --- | --- | --- | --- |
| Ig3 | 3.4 ± 0.7 | 4.0 ± 0.2 | 11.5 ± 2.2 |
| Ig3L | 2.0 ± 0.7 | 5.3 ± 0.5 | 20.2 ± 4.9 |
| Ig3-4 WT | 0.76 ± 0.18 | 3.9 ± 0.2 | 49.7 ± 7.1 |
| Ig3 + Ig4 | 3.93 ± 1.09 | 4.1 ± 0.40 | 14.0 ± 10.8 |
| Ig3L + Ig4 | 6.43 ± 1.79 | 6.0 ± 0.60 | 46.9 ± 23.1 |
| Ig3-4ΔSGHP | 4.4 ± 1.4 | 8.5 ± 0.7 | 89.5 ± 2.7 |
| Ig3-4ΔPHAR | 9.7 ± 1.6 | 9.4 ± 0.6 | 82.1 ± 8.3 |
| Ig3-4ΔHPHA | 10.7 ± 2.7 | 6.9 ± 0.7 | 84.0 ± 0.3 |
| Ig3-4Δ6 | 9.1 ± 1.5 | 8.2 ± 0.5 | 86.5 ± 2.5 |
| Ig3-4Δ21 | 15.7 ± 3.4 | 9.2 ± 0.9 | 82.1 ± 8.1 |
| Ig3-45L-4 | 12.6 ± 0.3 | 3.2 ± 0.5 | 6.9 ± 3.2 |
| Ig3-45L-5 | 2.5 ± 1.1 | 1.7 ± 0.2 | 10.7 ± 1.9 |
| RLinkerA | N/A | 0.9 ± 0.07 | 9.8 ± 11.8 |
| DELinkerA | 8.5 ± 4.6 | 6.1 ± 1.2 | 21.9 ± 7.9 |
| Scramble | 4.0 ± 1.3 | 7.0 ± 0.6 | 15.5 ± 7.3 |

### Deletions in linker region affect actin binding and bundling

Given the unique length of the linker region between the Ig3 and Ig4 domains of palladin, we sought to determine whether shorter versions would affect the actin binding and bundling. Therefore, we generated three mutants with short four residue deletions in the linker region covering different regions (Figure 3A) as well as two longer deletions of six and 21 residues (Figure 4A). Actin co-sedimentation assays demonstrated that all three of the short deletion mutants (ΔSGHP, ΔHPHA, and ΔPHAR) had decreased F-actin binding affinity (Table 1). Although all short deletion mutants maintained the same linker length, the deletion of different amino acid sets resulted in varying K_d_ values and binding curves (Figure 3B), despite some overlap in the amino acids deleted. Interestingly, while binding affinity was decreased, each mutant exhibited increased bundling efficiency at low concentrations of palladin (Figure 3C). Results from the two longer deletions in the linker (Figure 4B) showed that both constructs (Δ6 and Δ21) also had increased K_d_ values and reduced B_max_ values compared to wildtype (Table 1). Like the small deletion mutants, both Δ6 and Δ21 also displayed increased bundling efficiency at 5 μM (Figure 4C).

**Figure 3.**
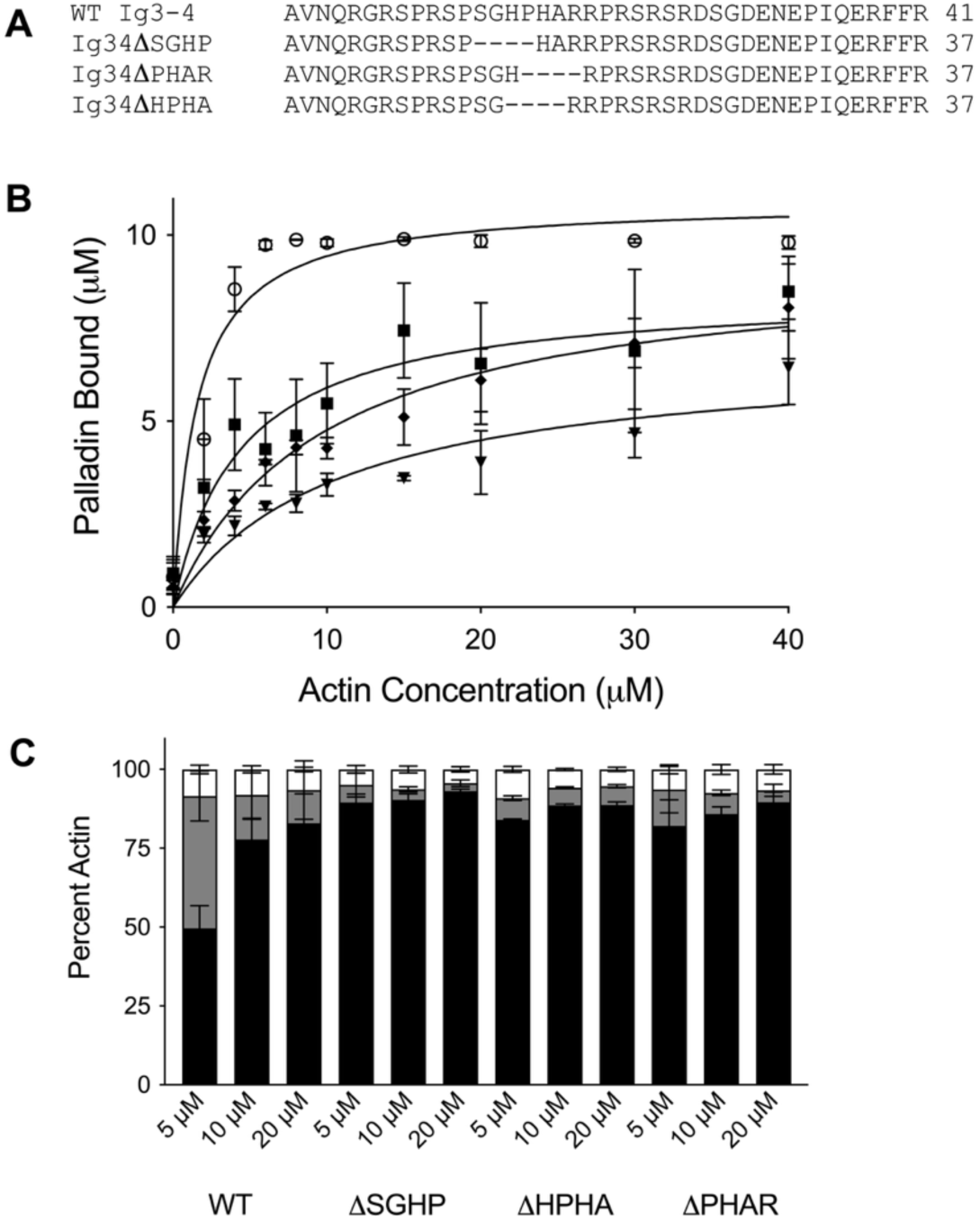
Small deletions in the Ig3–Ig4 linker reduce F-actin binding but increase bundling activity. **(A)** Sequence of the palladin Ig3–Ig4 linker region showing the positions of four-residue deletions in three mutant constructs: ΔSGHP, ΔPHAR, and ΔHPHA.**(B)** F-actin binding curves for wild-type Ig3–Ig4 (Ig3-4 WT, open circles), Ig3-4ΔSGHP (squares), Ig3-4ΔPHAR (diamonds), and Ig3-4ΔHPHA (triangles). Deletion mutants exhibit reduced binding affinity compared to wild-type, indicating that specific residues in the linker contribute to F-actin interaction. **(C)** F-actin bundling efficiency of wild-type and deletion constructs assessed by low-speed co-sedimentation. Bars represent the distribution of actin in the bundled fraction (black), pellet (gray), and supernatant (white). Deletion constructs show increased bundling efficiency relative to Ig3–4 WT. All data represent means ± standard deviation from at least three independent experiments.

**Figure 4.**
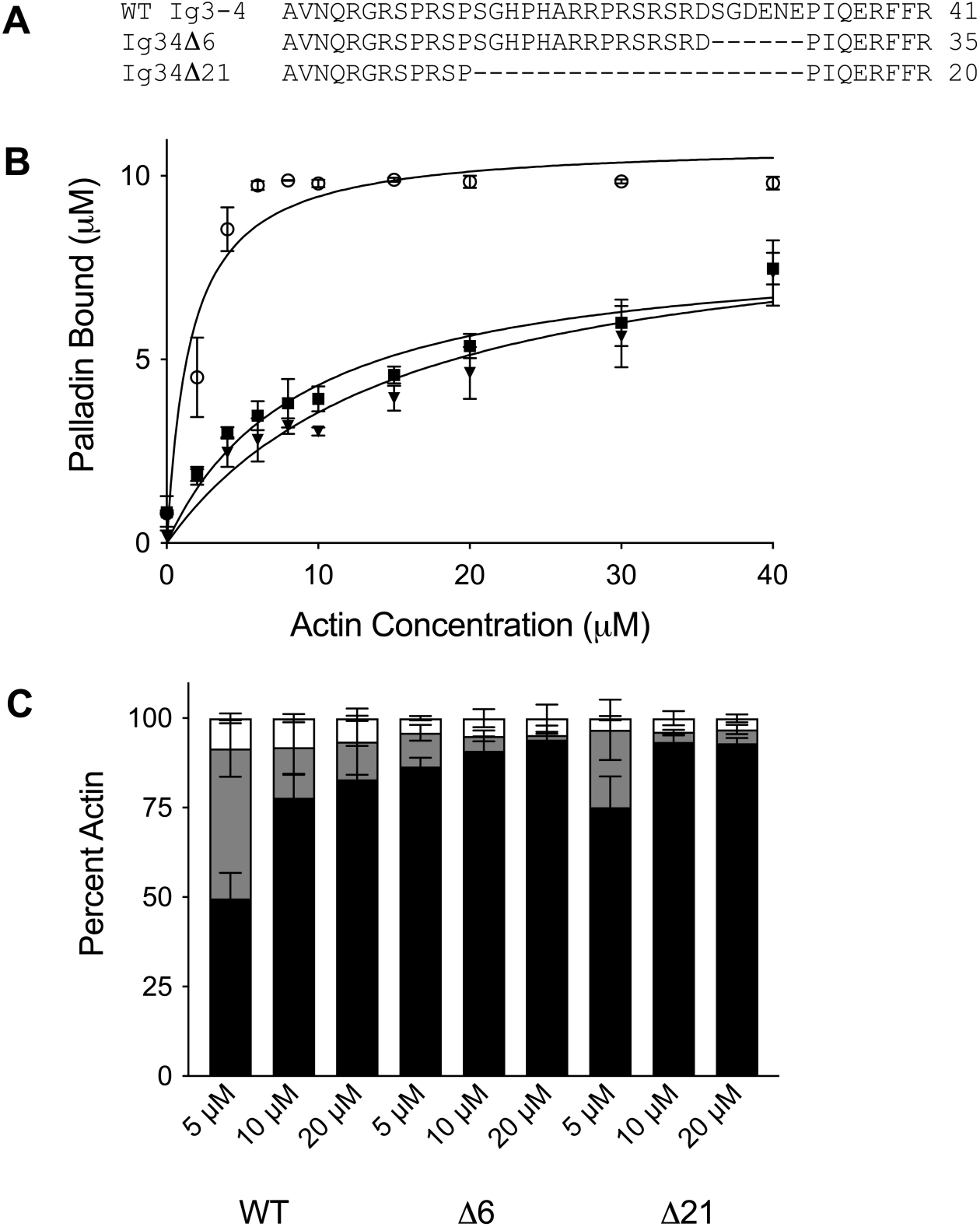
Longer deletions in the Ig3–Ig4 linker impair F-actin binding but not bundling. **(A)** Sequence of the palladin Ig3–Ig4 linker region indicating the positions and extents of two deletion constructs: Δ6 (six-residue deletion) and Δ21 (21-residue deletion). **(B)** F-actin binding curves for wild-type Ig3–Ig4 (Ig3-4WT, open circles), Ig3-4Δ6 (squares), and Ig3-4Δ21 (triangles). Both deletion constructs show reduced binding affinity and lower B_max_ values compared to WT, with the Δ21 mutant exhibiting the most severe impairment. **(C)** F-actin bundling efficiency of the constructs assessed via low-speed co-sedimentation. Bar graphs represent the proportion of actin found in the bundled fraction (black), pellet (gray), and supernatant (white). Longer deletion constructs show increased bundling efficiency relative to Ig3–4 WT. All data represent means ± standard deviation from at least three independent experiments.

### Ig4-Ig5 linker cannot compensate for the Ig3-Ig4 linker

To further investigate how linker length and sequence context influence palladin’s interaction with F-actin, we replaced the native 41-residue Ig3–Ig4 linker with the much shorter 7-residue linker that naturally connects Ig4 to Ig5 in full-length palladin (Figure 5A). This substitution was tested in two construct designs: one in which Ig3 was directly fused to Ig4 via the Ig4–Ig5 linker, and another in which Ig3 was fused to Ig5, bypassing Ig4 entirely.

**Figure 5.**
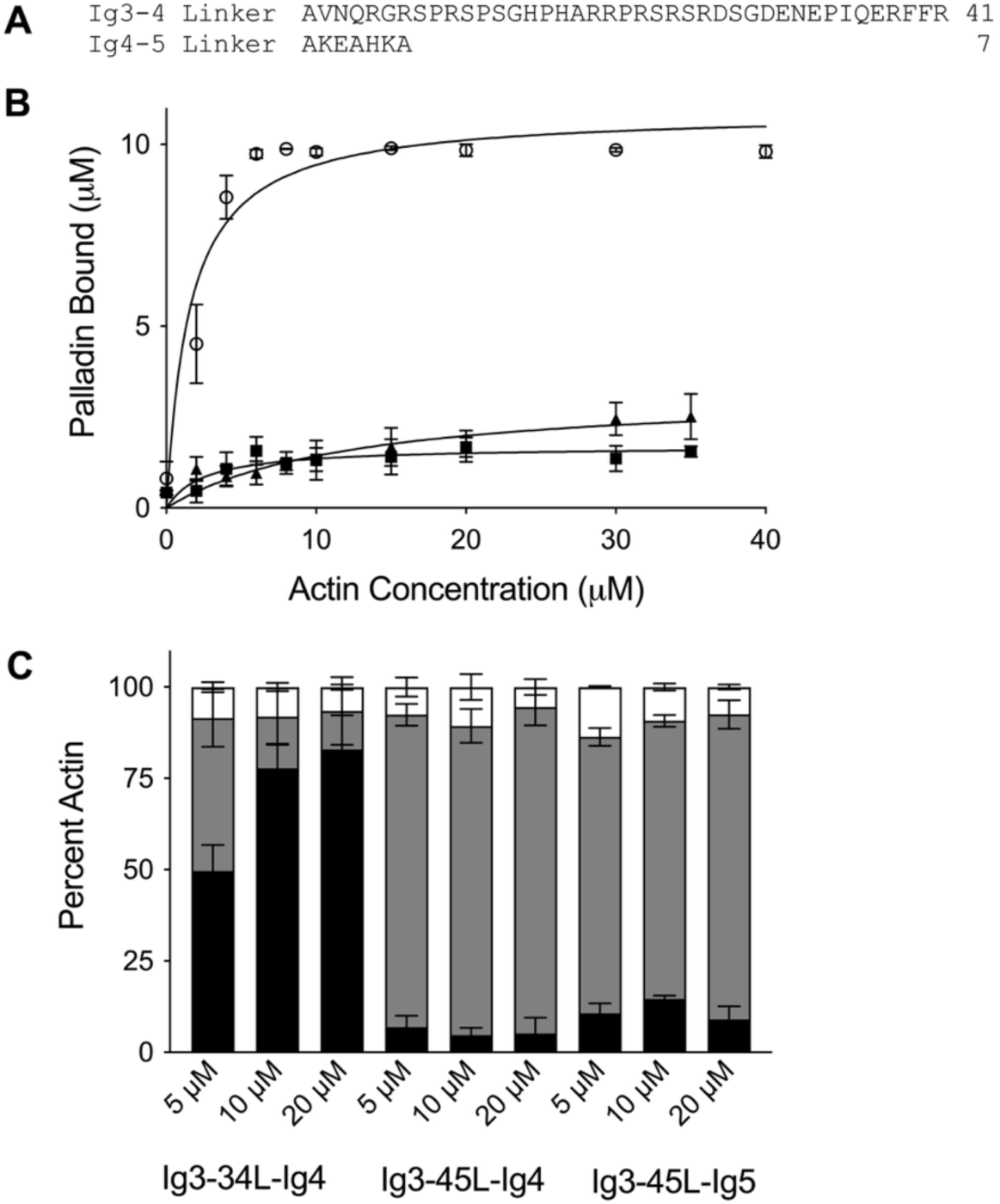
The Ig4–Ig5 linker and Ig5 domain cannot substitute for the native Ig3– Ig4 linker in F-actin binding and bundling. **(A)** Sequence overview comparing the native 41-residue Ig3–Ig4 linker region with the shorter 7-residue Ig4–Ig5 linker. **(B)** F-actin binding curves for wild-type Ig3–Ig4 (Ig3-4WT, open circles), Ig3–Ig4 connected by the Ig4–Ig5 linker (Ig3-45L-Ig4, triangles), and Ig3 connected to Ig5 via the Ig4–Ig5 linker (Ig3-45L-Ig5, squares). Both constructs with the substituted shorter linker exhibit decreased binding affinity and reduced B_max_ values compared to WT. **(C)** F-actin bundling efficiency assessed by low-speed co-sedimentation. Bar graphs indicate the distribution of actin in the bundled fraction (black), pellet (gray), and supernatant (white). Neither construct efficiently bundles actin, demonstrating that the shorter Ig4–Ig5 linker and the presence of Ig5 cannot compensate for the structural and functional role of the native Ig3–Ig4 linker. All data represent means ± standard deviation from at least three independent experiments.

Both constructs exhibited markedly impaired actin-binding properties, with significantly increased dissociation constants (K_d_) and reduced maximum binding capacities (B_max_) relative to the wild-type Ig3–Ig4 tandem construct (Figure 5B, Table 1). These results suggest that the shorter Ig4–Ig5 linker fails to provide the spatial flexibility or conformational freedom required for productive engagement of tandem domains with F-actin. Furthermore, neither construct was able to support efficient F-actin bundling (Figure 5C), consistent with their reduced binding affinity. This finding emphasizes that the linker’s role extends beyond merely connecting adjacent domains—it must also enable appropriate interdomain orientation and dynamic positioning necessary for multivalent interactions with actin filaments.

Interestingly, the construct connecting Ig3 directly to Ig5 also failed to recover bundling activity, indicating that simply increasing the number of Ig domains is insufficient to compensate for a suboptimal linker. The inability of either the short Ig4–Ig5 linker or a domain substitution to restore actin binding and bundling further underscores the unique functional role of the native Ig3–Ig4 linker. Its extended length, and possibly its intrinsic flexibility and electrostatic properties, appear essential for facilitating cooperative domain interactions and crosslinking of actin filaments. These findings support a model in which the length, flexibility, and specific composition of the Ig3–Ig4 linker are finely tuned to promote productive actin engagement. Replacing this region with a shorter linker likely imposes conformational constraints that prevent the two domains from adopting a geometry conducive to efficient F-actin binding and bundling.

### Overall charge and sequence order influence actin binding and bundling

To investigate how the composition and organization of the linker region influence actin interactions, we focused on its highly basic character—nearly 25% of the residues are arginine (R). We generated three Ig3-4 linker mutants to test how overall charge and sequence order affect F-actin binding and bundling. The first construct, RLinkerA, substituted all ten arginine residues in the linker with alanine (Figure 6A), effectively neutralizing its positive charge. Despite retaining an intact Ig3 actin-binding domain, this mutant exhibited a dramatic loss of both F-actin binding and bundling activity (Figure 6B and C). This result highlights the critical role of the linker’s positive charge in stabilizing interactions with the negatively charged actin filament surface, possibly by promoting electrostatic attraction or maintaining a favorable domain orientation for actin engagement.

**Figure 6.**
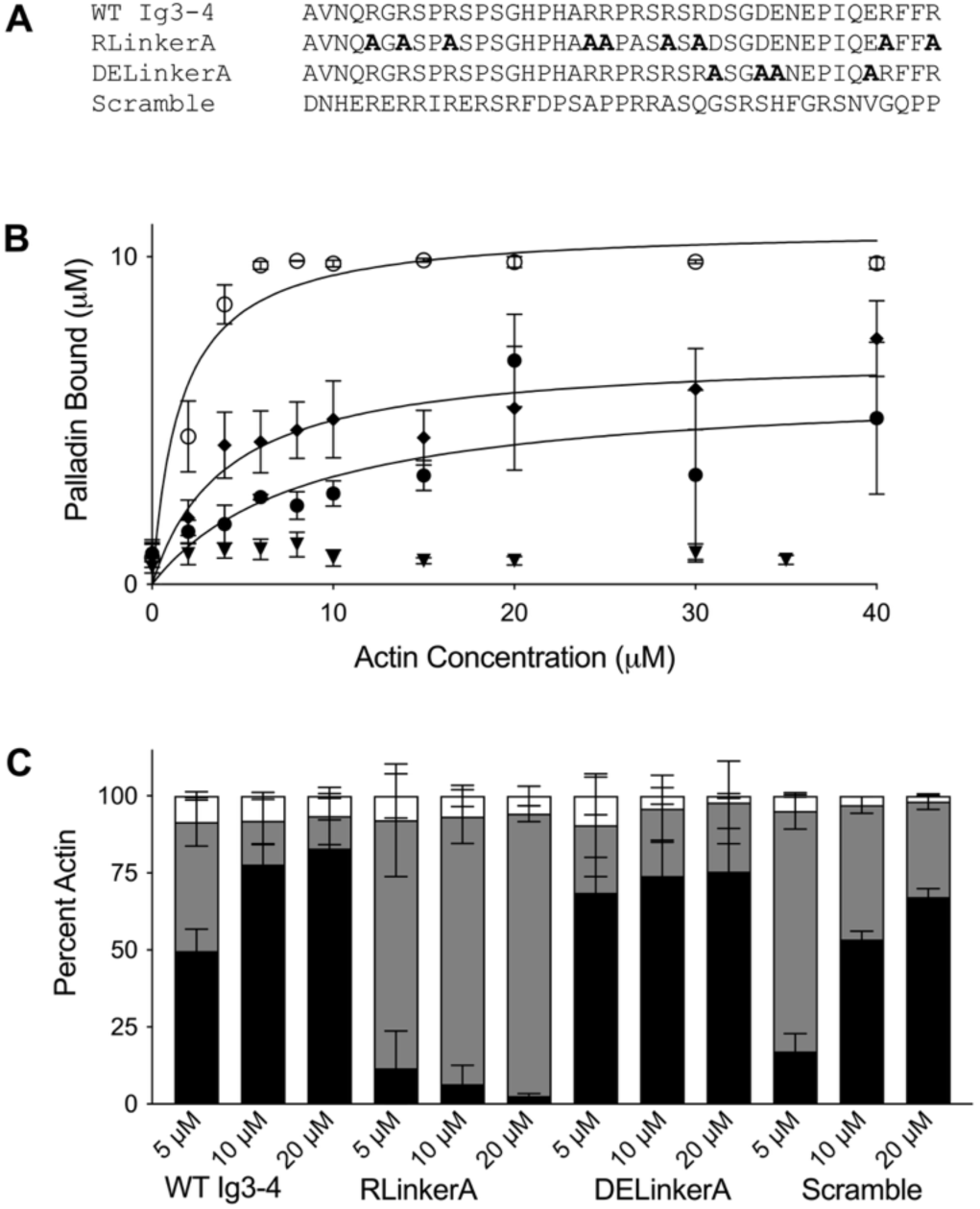
Linker charge and amino acid sequence order influence F-actin binding and bundling. **(A)** Sequence overview of the Ig3–Ig4 linker region highlighting mutated residues (in bold) in three constructs: RLinkerA (all nine arginines mutated to alanine), DELinkerA (four acidic residues neutralized), and LinkerScramble (sequence order randomized while preserving overall charge and length). **(B)** F-actin binding curves for wild-type Ig3–Ig4 (Ig3-4WT, open circles), RLinkerA (triangles), DELinkerA (squares), and LinkerScramble (diamonds). The RLinkerA mutant shows severely impaired actin binding, while DELinkerA exhibits reduced binding affinity and B_max_. LinkerScramble, despite retaining overall charge, also shows diminished binding, indicating that sequence order contributes to function. **(C)** F-actin bundling efficiency assessed via low-speed co-sedimentation. Bar graphs show the proportion of actin in the bundled fraction (black), pellet (gray), and supernatant (white). Loss of positive charge (RLinkerA) or disruption of sequence order (LinkerScramble) significantly reduces bundling efficiency, while DELinkerA enhances bundling at low concentrations despite reduced overall binding. All data represent means ± standard deviation from at least three independent experiments.

In contrast, the second construct, DELinkerA, neutralized all acidic residues within the linker, increasing its overall positive charge (Figure 6A). This modification resulted in a higher dissociation constant (K_d_) and reduced maximum binding (B_max_) compared to wild type (Table 1), suggesting that while increased positive charge may enhance initial filament recruitment, it may also disrupt optimal domain arrangement or binding specificity. Interestingly, DELinkerA promoted more efficient bundling at low concentrations (Figure 6C), potentially due to enhanced electrostatic bridging between adjacent filaments facilitated by the highly basic linker.

The third construct, LinkerScramble, maintained the native amino acid composition and net charge of the linker but randomized the residue order. This mutant displayed reduced actin-binding affinity and lower B_max_ compared to wild-type (Figure 6B), indicating that the spatial arrangement of charged and hydrophobic residues, not just their presence, is important for productive actin engagement. Moreover, LinkerScramble exhibited diminished bundling efficiency at all concentrations tested (Figure 6C), suggesting that specific sequence motifs or residue spacing within the linker may be required to stabilize inter-filament contacts or promote an extended conformation conducive to crosslinking.

Together, these data indicate that both the net charge and sequence order of the linker are critical for modulating palladin’s actin-binding and bundling activity. The positively charged linker likely plays a dual role: enhancing the local concentration of actin-binding domains near the filament surface and promoting crosslinking by bridging adjacent filaments. Disruption of either the electrostatic environment or sequence context impairs these functions, underscoring the linker’s active contribution beyond merely tethering adjacent domains.

### Linker mutations reduce thermal and chemical stability of Ig3–4 constructs

To ensure that the reduced actin-binding and bundling activities observed in several linker mutants were not due to gross structural perturbations, we first confirmed that their secondary structure content was comparable to wild-type using circular dichroism (CD) spectroscopy. DichroWeb results for WT and all mutants of palladin Ig3-4 maintained 48% beta-strand content, 4% alpha helix, and 48% loop or unstructured, which is consistent with the AlphaFold-predicted content.

To assess whether linker mutations altered the structural stability of palladin Ig3–4 constructs, we performed thermal denaturation monitored by circular dichroism (CD) and chemical denaturation using intrinsic tryptophan fluorescence. Wild-type Ig3–4 exhibited a melting temperature (T_m_) of 61.2 °C and a chemical denaturation midpoint (C_m_) of 3.98 M urea, consistent with a well-folded, stable tandem domain construct (Table 2, Figure 7A–B). In contrast, the Ig3–45L–Ig4 construct, in which the native 41-residue linker was replaced with the shorter Ig4–Ig5 linker, showed a modest but reproducible decrease in thermal stability (T_m_ = 56.95 °C), suggesting that linker shortening slightly destabilizes the tandem domain fold. Similarly, the RLinkerA and DELinkerA mutants—designed to neutralize or enhance the linker’s net charge— exhibited further reductions in thermal stability (T_m_ = 54.3 °C and 53.6 °C, respectively), indicating that electrostatic properties of the linker contribute to domain stability. The most dramatic destabilization was observed in the LinkerScramble mutant, which retained the native amino acid composition and charge but randomized residue order. This construct failed to yield a reliable thermal unfolding curve due to aggregation above 50 °C, precluding accurate T_m_ determination. However, the onset of aggregation and loss of CD signal at lower temperatures suggest a substantial decrease in thermal stability relative to WT.

**Figure 7.**
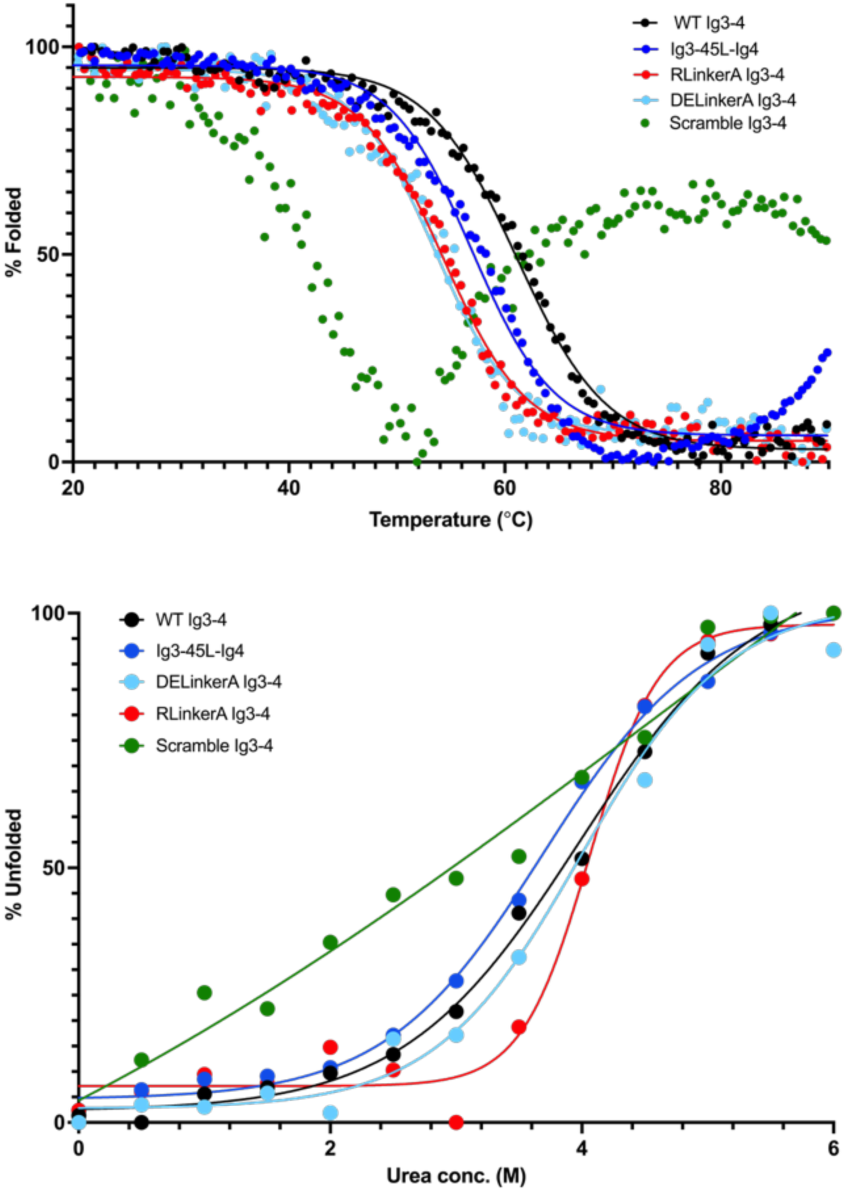
Linker mutations do not alter overall secondary structure or stability of Ig3–Ig4 constructs. **(A)** Circular dichroism (CD) thermal denaturation profiles for wild-type Ig3–Ig4 (WT, black) and linker mutants RLinkerA (red), DELinkerA (light blue), Ig3-45L-4 (dark blue), and LinkerScramble (green). CD measurements were performed at 206 nm, and unfolding transitions were monitored from 20–90 °C. Data were fitted to a two-state unfolding model to determine melting temperatures (T_m_), which are reported in Table 3. All constructs exhibit similar unfolding profiles and T_m_ values, indicating that linker mutations do not significantly impact the overall secondary structure or thermal stability except for the LinkerScramble mutant which unfolds at a significantly lower temperature and shows signs of aggregation at higher temperatures (>50 °C). **(B)** Intrinsic tryptophan fluorescence measurements of urea-induced unfolding show similar denaturation transitions for all constructs except for the LinkerScramble mutant. Emission intensity was normalized and fitted to a two-state denaturation model to calculate the midpoint of unfolding (C_m_) for each construct. C_m_ values are listed in Table 3. These results further support that most of the mutations do not perturb the native fold or stability of the Ig domains.

**Table 2:**
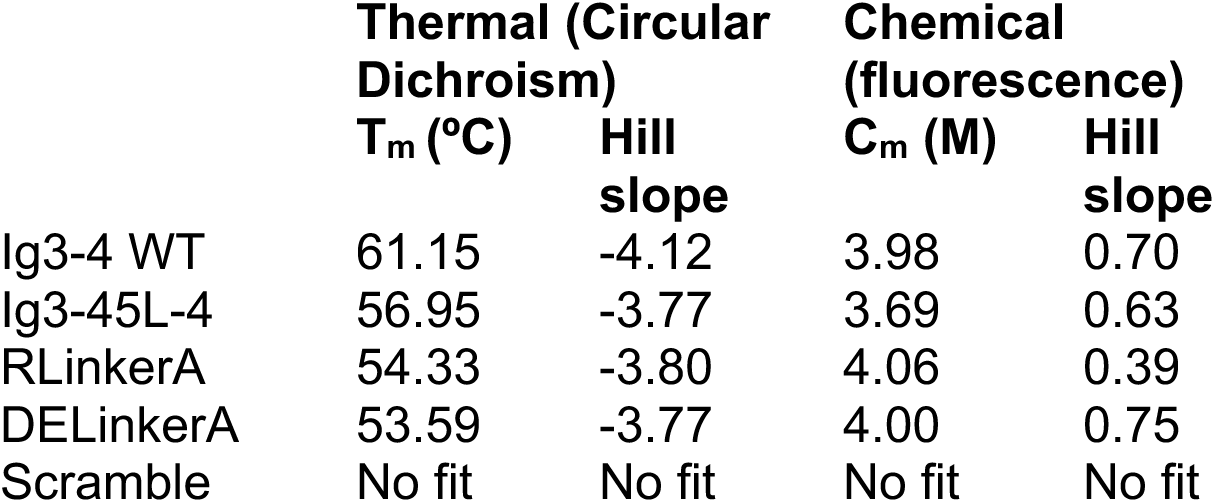
Unfolding parameters.

|  | **Thermal (Circular Dichroism)** |  | **Chemical (fluorescence)** |  |
| --- | --- | --- | --- | --- |
|  | **T<sub>m</sub> (°C)** | **Hill slope** | **C<sub>m</sub> (M)** | **Hill slope** |
| Ig3-4 WT | 61.15 | -4.12 | 3.98 | 0.70 |
| Ig3-45L-4 | 56.95 | -3.77 | 3.69 | 0.63 |
| RLinkerA | 54.33 | -3.80 | 4.06 | 0.39 |
| DELinkerA | 53.59 | -3.77 | 4.00 | 0.75 |
| Scramble | No fit | No fit | No fit | No fit |

Chemical denaturation profiles mirrored these trends. While WT Ig3–4 and the Ig3– 45L–Ig4, RLinkerA, and DELinkerA mutants all exhibited cooperative unfolding transitions with C_m_ values between 3.7–4.0 M urea, the LinkerScramble construct again failed to produce a reliable fit, consistent with misfolding or aggregation under denaturing conditions. Together, these results indicate that both the length and sequence composition of the Ig3–4 linker influence the structural stability of the tandem domain. While modest changes to linker length or charge reduce stability slightly, disruption of sequence order has a more severe impact, likely due to altered interdomain interactions or misfolding of the flexible linker region.

### Shorter linker region displays reduced conformational plasticity

Flexibility between domains in Ig3-4 has been previously offered as an explanation for actin-induced dimerization; however, this flexibility has not previously been assessed directly (16). To evaluate the impact of linker length on conformational plasticity, we collected SAXS data for Ig3-45L-Ig4 (with a seven-residue linker) and compared it with WT Ig3-4, Ig3, and Ig4. The normalized Kratky plot indicated peaks for both Ig3-45L-Ig4 and WT Ig3-4 that are consistent with multi-domain, flexible proteins, as indicated by the peak placement and dip in the curve around 0.3 Å^-1^ that does not return to baseline (Figure 8A). Small-angle X-ray scattering (SAXS) pairwise distance distribution [P(r)] analyses revealed distinct conformational characteristics for the isolated Ig domains and their linked constructs. The single Ig domains, Ig3 and Ig4, each exhibited a unimodal P(r) curve with a symmetric distribution and a single peak, consistent with the compact, globular nature of individual immunoglobulin-like folds (Figure 8B). In contrast, the Ig3–4 construct containing the native 41-residue linker displayed a broader P(r) curve with a pronounced trailing shoulder at higher interatomic distances. This feature suggests the presence of an extended conformation or significant flexibility between the two domains, likely due to the long unstructured linker that allows the domains to sample a wide range of relative positions. Conversely, the construct with a shortened seven-residue linker (Ig3-45L-4) showed a bimodal P(r) curve, with two distinct peaks. This pattern indicates a more restricted conformational ensemble in which the two domains are maintained in a closer, more fixed orientation, likely due to limited flexibility and enforced spatial proximity. Together, these SAXS data suggest that linker length and flexibility critically influence the relative arrangement and dynamics of tandem Ig domains, with implications for their structural organization and potential interaction interfaces.

**Figure 8.**
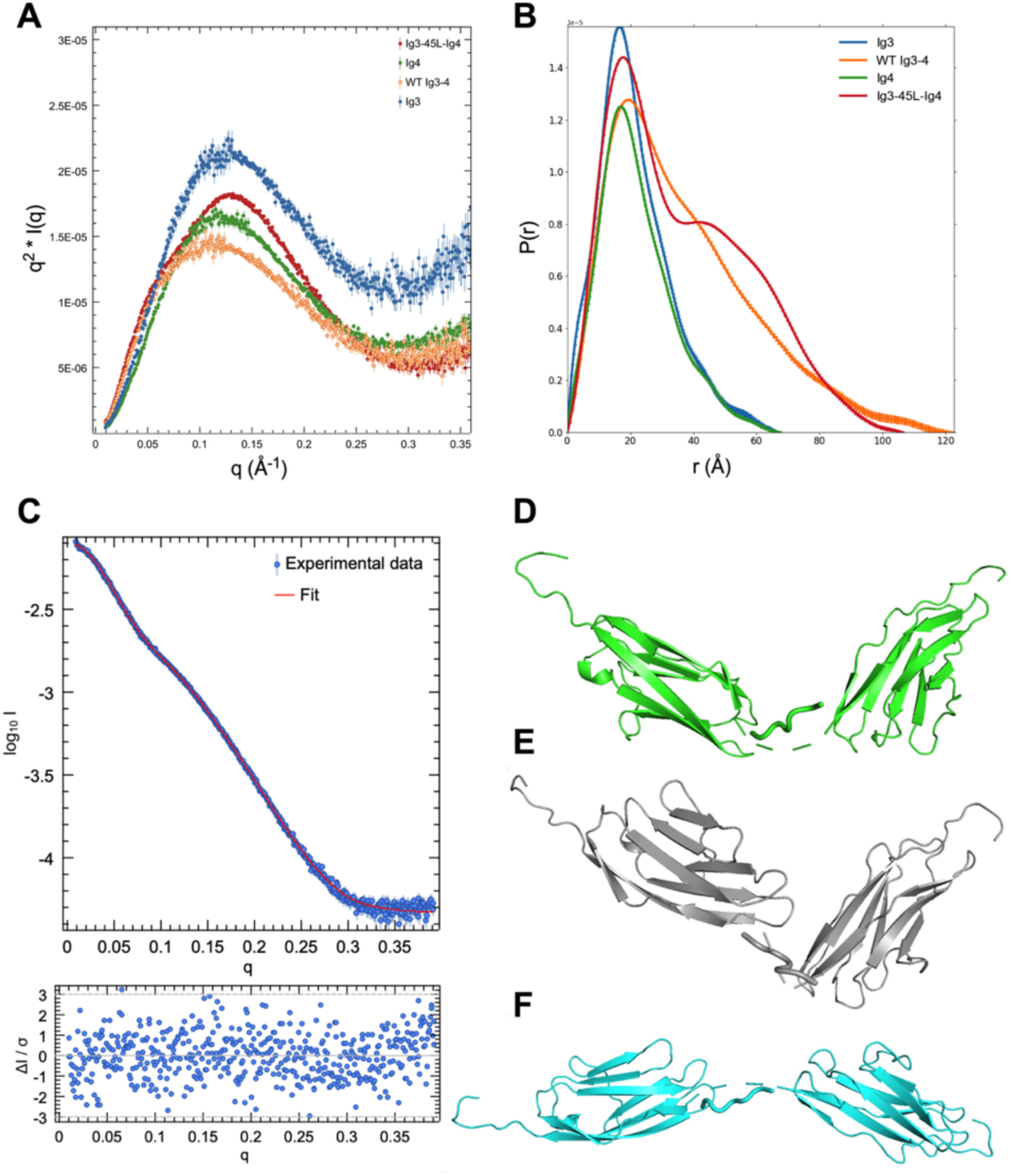
SAXS analysis reveals differences in domain organization and flexibility among palladin constructs. **(A)** Kratky plots for individual and tandem Ig domains: Ig3 (blue), Ig4 (green), Ig3–4 (orange), and Ig3-45L-Ig4 (red). Ig3 and Ig4 show characteristic bell-shaped curves consistent with compact, folded domains, while the tandem constructs display broader or shifted profiles indicative of increased flexibility or extended conformations. **(B)** Pair-distance distribution functions (P(r)) for the same constructs. Ig3 and Ig4 exhibit single, symmetric peaks consistent with globular shapes. **(C)** Experimental SAXS scattering profiles (blue) are well-fitted by theoretical curves generated from EOM-derived ensembles (red), with residuals plotted below (𝜒^2^ = 1.19). The ensemble includes three conformers that represent the structural heterogeneity observed in solution (see **D–F**).

To investigate domain orientations and linker flexibility, structural models were evaluated against SAXS data using both rigid-body and ensemble-based approaches. CRYSOL, a SAXS-based rigid-body modeling tool, was used to optimize the relative positions and orientations of the known high-resolution structures of the Ig domains, while approximating the missing linker region (17). This fitting yielded a 𝜒² value of 3.08, indicating some mismatch between the model and experimental data, likely due to conformational heterogeneity not captured by a single static structure. (See Supplemental Figure S1 for Guinier plot and CORAL analysis).

To model this flexibility in the shortened linker variant Ig3-45L-4, we employed the Ensemble Optimization Method (EOM) (18, 19). A large pool of random conformers was generated using RANCH, based on the known Ig domain structures, and GAJOE applied a genetic algorithm to select sub-ensembles that best fit the experimental scattering data. The selected ensemble produced theoretical scattering curves in good agreement with the experimental profiles (𝜒² value of 1.19), supporting the presence of a dynamic conformational ensemble in solution. The experimental scattering profile was well reproduced by the selected ensemble, as demonstrated by the close agreement between the experimental and calculated curves in Figure 8C. This ensemble consisted of three representative conformers capturing the range of observed structural variability (Figure 8D–F).

Compared to the previously published EOM analysis of wild-type Ig3-4 [17], the Ig3-45L-4 variant exhibited a narrower range of conformations, with R_g_ values spanning 25.75–33.92 Å and D_max_ values from 95.01–121.81 Å. In contrast, the wild-type ensemble displayed a broader range of structural variability, with R_g_ values between 23.55–38.27 Å and D_max_ values of 79.15–115.95 Å. These results suggest that the shortened linker in Ig3-45L-4 constrains the molecule to a more extended but less conformationally diverse ensemble, likely due to reduced interdomain flexibility. Conversely, the longer linker in wild-type Ig3-4 enables a wider range of interdomain motions, allowing both compact and extended conformations. This increased flexibility may facilitate dynamic functional interactions or structural rearrangements not accessible to the shortened variant.

## Discussions

Our findings demonstrate that the linker connecting the Ig3 and Ig4 domains of palladin is a critical determinant of its ability to bind and bundle F-actin. While prior studies have shown that the tandem Ig3–4 domains bind F-actin with greater affinity than the individual Ig3 domains, neither the isolated Ig4 domain nor the linker alone exhibit binding activity (14, 16). Nevertheless, the specific functional contribution of the intervening linker between these domains has remained largely uncharacterized. Here, we show that this linker is not merely a passive connector but an active modulator of interdomain flexibility, actin engagement, and filament crosslinking.

### Linker region enhances actin-binding and bundling cooperativity

Consistent with earlier reports that Ig3 alone has variable bundling capacity depending on conditions and concentration (14, 15), our data confirm that Ig3 can mediate weak actin binding and limited bundling. However, inclusion of the native linker (Ig3L) significantly enhances both activities, even in the absence of Ig4. These results suggest that the linker promotes a more favorable spatial orientation of the actin-binding domain relative to the filament surface—an effect that becomes more pronounced in the WT tandem Ig3–4 construct. This enhancement is lost when Ig4 is added in trans, highlighting the requirement for covalent connectivity and structural integration. Similar domain cooperation has been described in other actin-binding proteins, such as filamin and spectrin, where flexible linkers regulate multivalent interactions and actin crosslinking capacity (20, 21).

### Linker deletions uncouple binding from bundling

Surprisingly, small deletions within the linker reduced actin-binding affinity but increased bundling activity at low protein concentrations. One explanation is that the shortened linker restricts domain flexibility, stabilizing an extended domain configuration better suited to spanning and crosslinking adjacent filaments. This mirrors observations where shorter or stiffer crosslinkers such as α-actinin bind actin with high affinity yet form less extensible bundles (22), whereas flexible or longer linkers (e.g. filamin) support bundling through wider spacing and cooperative crosslinking, often with lower individual-domain affinity (23, 24). These results imply that optimal bundling does not always correlate with maximal binding affinity, and that structural constraints imposed by the linker may selectively favor crosslinking geometries.

### The Ig4–Ig5 linker cannot substitute for Ig3–Ig4

Our finding that the shorter Ig4–Ig5 linker, with only seven residues, fails to restore activity when substituted between the Ig3–4 domains further supports the specialized role of the native linker. Despite the conserved fold of immunoglobulin domains, the linkers connecting them vary widely in length and sequence (Figure 1A). In palladin, the 41-residue Ig3–Ig4 linker appears uniquely adapted to balance flexibility and spacing, enabling cooperative engagement of actin filaments. Similar specificity in linker function has been noted in other modular actin-binding proteins, including titin and α-actinin, where linker constraints modulate domain mobility and interaction potential (25, 26).

### Electrostatic tuning and sequence order are critical

Charge-based tuning of actin-binding domains is a well-documented mechanism for modulating filament interactions (27-31). Here, we show that neutralizing basic residues in the linker abolishes actin binding and bundling, while increasing net positive charge paradoxically reduces binding affinity but enhances bundling. These effects likely reflect changes in electrostatic steering, domain orientation, or filament bridging potential. Moreover, scrambling the linker sequence—without altering its net charge—still impairs function, indicating that spatial arrangement of residues is essential for productive interaction. This finding is consistent with studies on intrinsically disordered regions (IDRs) in actin regulators like coronins (32) and WASP (33), where residue patterning influences conformational sampling and target engagement.

### Linker composition modulates structural stability of tandem Ig domains

Our biophysical analyses reveal that the structural stability of the palladin Ig3–4 tandem domain is sensitive to both the length and sequence composition of the interdomain linker. Wild-type Ig3–4 exhibits robust thermal and chemical stability, consistent with a well-folded tandem domain architecture. However, targeted mutations within the linker region—designed to alter its length, charge, or sequence order—resulted in varying degrees of destabilization, highlighting the importance of this region in maintaining domain integrity.

Substitution of the native 41-residue linker with the shorter Ig4–Ig5 linker (Ig3–45L–Ig4) led to a modest but reproducible decrease in thermal stability (T_m_ = 56.95 °C), suggesting that linker shortening imposes conformational constraints that subtly destabilize the tandem fold. More pronounced effects were observed for the RLinkerA and DELinkerA mutants, which respectively neutralize or enhance the linker’s net positive charge. Both constructs exhibited reduced thermal stability (T_m_ = 54.3 °C and 53.6 °C, respectively), indicating that electrostatic interactions within the linker contribute to the overall stability of the domain pair. These findings are consistent with prior studies showing that charged residues in flexible regions can influence protein folding landscapes and interdomain packing.

The most striking destabilization was observed in the LinkerScramble mutant, which retains the native amino acid composition and net charge but disrupts the sequence order. This construct aggregated at temperatures above 50 °C and failed to yield a reliable thermal unfolding curve, suggesting a substantial loss of structural integrity. The inability of this mutant to maintain a stable fold—despite preserving overall charge— underscores the importance of residue patterning and sequence context in disordered linker regions. These results align with emerging evidence that sequence order within intrinsically disordered regions (IDRs) can dictate conformational sampling, folding cooperativity, and interaction specificity.

Chemical denaturation profiles mirrored the thermal trends, with WT and most mutants exhibiting cooperative unfolding transitions, while the LinkerScramble construct again failed to produce a reliable fit. Together, these data suggest that the Ig3–4 linker contributes not only to actin-binding and bundling activity but also to the structural stability of the tandem domain. Disruption of linker properties—whether by shortening, charge alteration, or sequence scrambling—can compromise both functional and structural integrity, reinforcing the view that this region is a finely tuned module essential for palladin’s cytoskeletal role.

### Structural flexibility governs tandem domain behavior

SAXS-based modeling provides further insight into how linker properties modulate domain architecture. The wild-type Ig3–4 tandem adopts a dynamic, flexible ensemble with broad R_g_ and D_max_ distributions, consistent with an unstructured linker that allows extensive conformational sampling. In contrast, shortening the linker constrains the ensemble to fewer, more extended conformations. These observations are in line with prior SAXS and NMR studies showing that interdomain linkers regulate the range of accessible conformations and affect ligand binding and allostery (34-36). Notably, the decreased conformational plasticity of the Ig3-45L-4 construct may explain its reduced ability to adopt binding-competent orientations on the actin filament surface.

### Implications for palladin function and cytoskeletal regulation

Together, our data support a model in which the Ig3–Ig4 linker acts as a tunable element that coordinates spatial arrangement, domain flexibility, and electrostatic compatibility to mediate F-actin engagement and crosslinking. The failure of non-native linkers or domain swaps to restore activity underscores the evolutionary specialization of this region. These results have broader implications for the regulation of actin-binding proteins with modular architectures. In particular, linker-mediated modulation of domain flexibility may represent a general strategy for tuning cytoskeletal organization, mechanosensing, and signaling responses (37). Further exploration of the *in vivo* relevance of linker mutations—and potential regulation by phosphorylation or alternative splicing—may provide insight into palladin’s roles in cell migration, adhesion, and cancer metastasis (38, 39). Overall, our study highlights the importance of linker regions in shaping the functional landscape of actin-binding proteins.

## Materials and Methods

### Mutagenesis

Targeted mutagenesis of the linker region between the Ig3 and Ig4 domains of palladin was performed using the Q5® Site-Directed Mutagenesis Kit (New England Biolabs, Cat. #E0554S), following the manufacturer’s protocol. Mutagenic primers were designed using the NEBaseChanger tool (https://nebasechanger.neb.com/) to introduce specific amino acid substitutions, deletions, or insertions within the linker sequence. Primers were synthesized by Integrated DNA Technologies (IDT) and resuspended to a final concentration of 10 µM. Mutations were introduced into the Ig3-4 linker region using the Q5® Site-Directed Mutagenesis Kit (NEB) following the manufacturer’s protocol. PCR amplification with mutagenic primers was performed according to kit specifications, followed by KLD (kinase, ligase, DpnI) treatment and transformation into *E. coli* DH5α cells. Plasmid DNA from individual colonies was sequenced to confirm the presence of desired mutations.

### Palladin Protein Expression and Purification

Palld-Ig3 and Palld-Ig4 domains in the pTBSG expression vector as well as Palld-Ig3-4 and all its mutant forms in the pTBMalE expression vector were overexpressed in BL21 (DE3) *Escherichia coli* cells cultured in autoinduction media as previously described (15). Cell suspensions were lysed by sonication and centrifuged to pellet cell debris. Supernatant was purified via PROTEINDEX™ HiBond™ Ni-NTA Agarose according to manufacturer’s guidelines. After loading the supernatant, column was washed with Wash Buffer (20 mM Na_2_HPO_4_ pH 7.4, 100 mM NaCl, 5 mM imidazole) and proteins were collected using the Elution Buffer (20 mM Na_2_HPO_4_ pH 7.4, 100 mM NaCl, 250 mM imidazole). TEV protease was used to cleave the N-terminal tag (His_6_ for pTBSG constructs and His_6_-MBP for pTBMalE constructs). For constructs in the pTBMalE vector, the MBP tag was separated from the digested protein by using Amylose Resin (New England Biolabs) in a gravity flow column according to manufacturer instructions. Lastly, cation exchange chromatography (SP sepharose, GE Healthcare Life Sciences) was used to isolate the target protein. Pure fractions were confirmed by SDS-PAGE and dialyzed to storage buffer at 4 °C (20 mM HEPES, pH 7.4, 1 mM DTT, 100 mM NaCl).

### Actin Co-sedimentation

Actin was purified from rabbit muscle acetone powder purchased from PelFreeze Biologicals using a method adapted from Spudich and Watt (40). Binding assay samples contained 10 μM actin binding protein and increasing concentrations of polymerized actin (0-40 μM). F-buffer (10 mM Tris/HCl pH 7.5, 100 mM KCl, 2 mM MgCl_2_, 2 mM DTT) was added to each tube for a final reaction volume of 100 μL. Samples were made in triplicate and incubated at room temperature for 30 minutes then centrifuged at 150,000 x*g* using the Ti-45 rotor in a Beckman Coulter Optima LE-80K Ultracentrifuge for 30 minutes at 4 °C. Supernatants were carefully separated. Pellets were resuspended in 100 μL of 2X Tris-glycine electrophoresis buffer (50 mM Tris pH 8.3, 500 mM glycine, 0.2% SDS). All fractions were separated by SDS-PAGE. Resulting gels were stained with Coomassie Brilliant Blue and scanned using an Epson WorkForce WF-3620 scanner for quantification of protein bands using ImageJ software (41).

Bundling assay samples contained 10 μM polymerized actin and 0, 5, 10, or 20 μM purified actin binding protein. An additional control containing only 10 μM purified protein (no actin) was also used. F-buffer was added to each tube for a final volume of 100 μL followed by incubation at room temperature for 30 minutes. Samples were centrifuged at 5,000 x*g* for 10 minutes at room temperature on a Thermo-Scientific Sorvall Legend Micro 21 Centrifuge. Supernatants were separated and sedimented protein (bundles) was resuspended in 100 μL of 2X Tris-glycine electrophoresis buffer. Supernatants from the slow spin step were ultracentrifuged at 150,000 x*g* using Ti-45 rotor in a Beckman Coulter Optima LE-80K Ultracentrifuge for 30 minutes at 4 °C. Supernatants from this fast spin step were separated and pellets were resuspended in 2X Tris-glycine electrophoresis buffer. All fractions were separated by SDS-PAGE. Gels were scanned and analyzed in the same way as the binding assay gels except actin bands were quantified.

### Circular dichroism

Purified palladin samples were extensively dialyzed into CD buffer (10 mM phosphate buffer, 50 mM sodium fluoride, 1 mM TCEP, pH 7.5). Spectra were recorded on a Jasco J810 spectropolarimeter at 30 μM, 15 μM, and 10 μM of each protein between 260 to 185 nm. Voltage was monitored during all scans to ensure it did not exceed 800 Hz. Thermal denaturation was monitored on 15 μM protein at 206 nm in a 0.1 mm water-jacketed cell that was heated from room temperature to 90 *°* C as controlled by a JULABO water circulator. This wavelength was chosen because it showed the greatest difference between the native and unfolded samples in all cases. Melting temperatures (T_m_) and Hill coefficients were determined by nonlinear fitting of the thermal unfolding curves using a Boltzmann sigmoidal curve in GraphPad Prism®10.

CD spectra were converted to mean residue ellipticity and HT voltage, then analyzed using DichroWeb (42). Scans were input with a wavelength range of 260–185 nm; the lowest wavelength used was adjusted based on voltage quality. Secondary structure content (*α* -helix, *β* -sheet, and random coil) was determined using the K2D analysis method. All output graphs and tables were saved for comparison.

### Fluorescence spectroscopy

Purified proteins (WT Ig3-4 and various mutants) were diluted to a final concentration of 10 μM in 100 mM phosphate buffer (pH 7.5) with varying urea concentrations (0-6 M urea). Samples were incubated overnight at room temperature and fluorescence emission spectra were collected on a PTI spectrofluorometer with excitation at 275 nm and emission observed at 280-400 nm. The fraction folded was calculated from emission spectra peak maxima at 350 nm and plotted against urea concentration. The normalized data were fitted to a Boltzmann sigmoidal curve in GraphPad Prism®10 software to determine the midpoint of denaturation (C_M_) and Hill Slope values.

### Small Angle X-ray Scattering

After dialysis of purified palladin domains into HEPES storage buffer, proteins were further purified by size exclusion chromatography on the Cytiva Superdex 200 10/300 GL column in preparation for SAXS data collection. Ig3 and Ig4 were concentrated and prepared directly for SAXS, while Ig3-4 and Ig3-45L-4 were subjected to a final purification step on the Cytiva HiLoad 16/600 Superdex 200 pg column to remove the remaining contaminants. Concentration series were prepared for each protein: 0.5-4.3 mg/mL for Ig3, 0.5-10 mg/mL for Ig4, 0.5-7.4 mg/mL for Ig3-4 and 0.5-10 mg/mL for Ig3-45L-4 by dilution with storage buffer. SAXS data was collected at twelve frames per sample with 1 second exposures at the Stanford Synchrotron Radiation Library (SSRL) (43). Resulting data was automatically buffer-subtracted and analyzed on-line with SAXSPipe (44). The ATSAS software package, including PRIMUS, CRYSOL and SASREF, was used for further analysis (45, 46). Buffer-subtracted files were compared at every concentration for each protein. Minimal concentration dependence was observed for all domains, so the low q range of a low concentration sample was merged with the high q range of a higher concentration sample using PRIMUS software (45). PRIMUS was further used for each merged data curve to estimate radius of gyration (R_g_) and the forward scattering intensity I(0) from the Guinier plot and Porod volume and R_max_ from the distance distribution analysis.

Prior to structural modeling, sequence and structure files were optimized as follows. For Ig3, the N-terminal “SNA” and “GGS” residues were removed from the sequence and PDB files, respectively, for sequence continuity. For Ig4, the C-terminal “AHK” and “SGPSSG” residues were removed from the sequence and PDB files, respectively.’ Furthermore, the sequence was modified with I179V and N224S to match the published structure.

CRYSOL (maximum *s* value 0.4) (46, 47) and SASREF (17) were used to check data quality for Ig3 and Ig4 with known structures, while the CORAL2.8.2 (48) was used to create and compare theoretical scattering curves to the experimental data and to create an initial rigid-body model. The AlphaFold predicted structure for Ig3-45L-Ig4 was used for experimental data comparison with CORAL rigid body modeling (48). SASREF was run with the Ig3-45L-Ig4 experimental data and amplitudes for the individual domains were computed by CRYSOL.

The molecular ensemble model was generated by the Ensemble Optimization Modelling (EOM) software (18, 19), a pipeline of SAXS analysis programs. Initially, the RANCH package was used to generate a pool of 10,000 models based on two rigid bodies defined by the known protein structures (PDB: 2LQR and 2DM3) while the missing linker region (residues 97-104 in the tandem Ig3-45L-Ig4) was allowed to move freely. A theoretical scattering intensity curve based on the ensemble was generated using GAJOE, which fits the experimental SAXS data with theoretical scattering curves generated from weighted-average sub-ensembles. Since GAJOE is a genetic algorithm that produces degenerate results across independent trials, we conducted five separate runs and assessed the solutions for convergence. Default parameters were used in all cases. EOM is semi-quantitative approach to analyze the flexibility and size distribution of possible configurations and we were able to obtain optimized ensembles with a fit to the experimental scattering data (𝜒^2^ ∼1.187) (Supplemental Figure S1 for Guinier plot and Figure S2 CORAL analysis).

## Data Availability

SAXS data for Ig3-45L-Ig4 has been deposited in the SAXBDB (SASDX77). Authors agree to make any materials, data, and associated protocols available upon request.

## CRediT Author Contribution

M.R.B. conceived and supervised the project. M.R.B., R.A.S., R.V., J.G.B. and M.J.L. provided the resources. M.R.B. and R.A.S. designed the experiments. R.A.S., C.W.B., L.M.H., and N.H.T. performed the experiments and analyzed the data. M.R.B. and R.A.S. prepared the manuscript. All authors reviewed and edited the manuscript.

## Acknowledgements

We thank Drs. Thomas Weiss (SSRL, USA) and Allyn Schoeffler (Loyola University New Orleans, USA) for help with SAXS data collection and analysis support. The FP7 WeNMR (project# 261572), H2020 West-Life (project# 675858), the EOSC-hub (project# 777536) and the EGI-ACE (project# 101017567) European e-Infrastructure projects are acknowledged for the use of their web portals, which make use of the EGI infrastructure with the dedicated support of CESNET-MCC, INFN-LNL-2, NCG-INGRID-PT, TW-NCHC, CESGA, IFCA-LCG2, UA-BITP, TR-FC1-ULAKBIM, CSTCLOUD-EGI, IN2P3-CPPM, CIRMMP, SURFsara and NIKHEF, and the additional support of the national GRID Initiatives of Belgium, France, Italy, Germany, the Netherlands, Poland, Portugal, Spain, UK, Taiwan and the US Open Science Grid.

## Abbreviations

A: alanine
Akt1: oncogene for kinase with role in cell growth, division and survival
B_max_: maximum specific binding capacity
CD: circular dichroism
C_m_: denaturation midpoint concentration
D: aspartic acid
D_max_: maximum linear dimension of particle in SAXS sample
DTT: dithiothreitol
E: glutamic acid
EOM: ensemble optimization method
F-actin: filamentous or polymerized form of actin
G-actin: globular or monomeric actin
IDRs: intrinsically disordered regions of proteins
Ig: immunoglobulin
K_d_: dissociation constant
NMR: nuclear magnetic resonance
PBD: protein database
P(r): pair-wise distance distribution function
PTMs: post-translational protein modifications
R: arginine
R_g_: radius of gyration
SAXS: small angle X-ray scattering
SDS-PAGE: sodium dodecyl sulfate polyacrylamide gel electrophoresis
TEV: Tobacco Etch Virus
T_m_: thermal denaturation midpoint temperature
WASP: Wiskott-Aldrich syndrome protein
χ^2^: chi-squared statistical test to compare observed with expected data

