## Supplemental for "The Role of Linker Length and Composition in Actin Binding and Bundling by Palladin"

**Running title:** Palladin Linker Modulates Tandem Domain Activity

Contents

1 Supplementary Figures

a. Figure S1 – Guinier region linear fit for Ig3-45L-Ig4

b. Figure S2 – CORAL fit for Ig3-45L-Ig4

2 Supplementary Notes

a. Supplementary Note 1. Detailed sequence information of the protein constructs

b. Supplementary Note 2. Primer sequences used for all mutations.

c. Supplementary Note 3. SAXS data deposition link

Supplemental Materials

Supplemental Figure S1

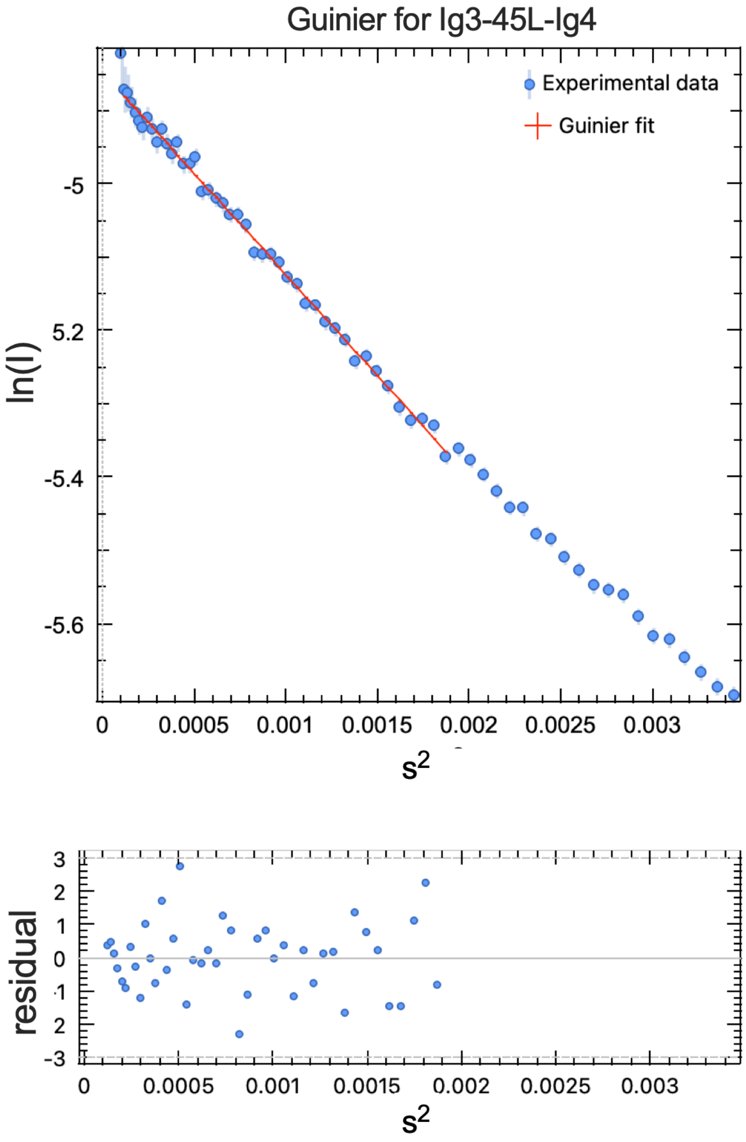


**Figure S1. Guinier region and linear fit with residuals for Ig3-45L-Ig4.** Experimental SAXS data in blue shown with Guinier linear fit region (red). Residuals are shown below.

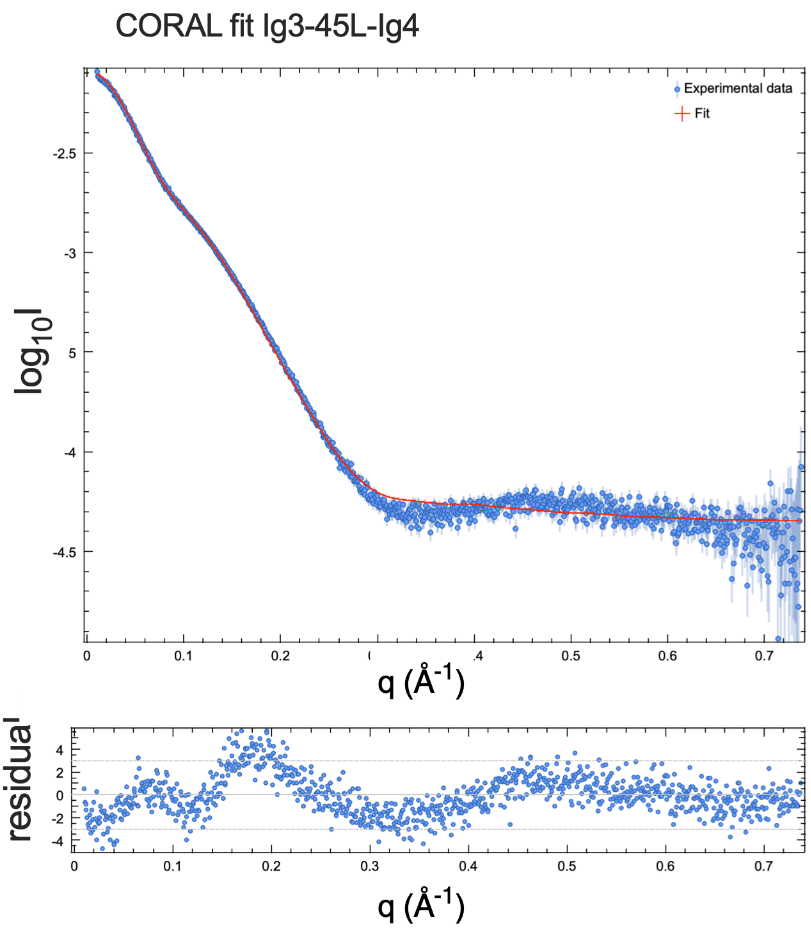


**Figure S2. CORAL (Complexes with Random Loops) SAXS-based rigid body modeling of Ig3-45L-Ig4.** SAXS scattering curve of palladin Ig3-45L-Ig4 shown in blue with CORAL rigid-body and linker modeling fitting to experimental SAXS data (red line). Residuals are shown below. This fitting yielded a $\chi^2$ value of 3.08, indicating some mismatch between the model and experimental data, likely due to conformational heterogeneity not captured by a single static structure.

Supplemental Materials

**Supplemental Note 1:** Protein sequences
Legend:
red = plasmid, green = linker, **bold** = mutations, blue = domain swap

Wildtype Sequences

- Ig3 WT

SNANATAPFFEMKLKHYKIFEGMPVTFTCRVAGNPKPKIYWFKDGKQISPKSDHYTIQRDLDGTCSLH
TTASTLDDDGNYTIMAANPQGRVSCTGRLMVQAVNQRGRS

- Ig4 WT

SNATVQERFFRPHFLQAPGDLTVQEGKLCRMDCKVSGLPTPDLSWQLDGKPIRPDSAHKMLVRENGVH
SLIIEPVTSRDAGIYTCIATNRAGQNSFNLELVVAAKEAHK

- Ig3-4WT

SNANATAPFFEMKLKHYKIFEGMPVTFTCRVAGNPKPKIYWFKDGKQISPKSDHYTIQRDLDGTCSLH
TTASTLDDDGNYTIMAANPQGRVSCTGRLMVQAVNQRGRSPRSPSGHPHARRPRSRSRDSGDENEPIQ
ERFFRPHFLQAPGDLTVQEGKLCRMDCKVSGLPTPDLSWQLDGKPIRPDSAHKMLVRENGVHSLIIEP
VTSRDAGIYTCIATNRAGQNSFNLELVVAAKEAHK

Mutant Sequences

- Ig3-4Δ21

SNANATAPFFEMKLKHYKIFEGMPVTFTCRVAGNPKPKIYWFKDGKQISPKSDHYTIQRDLDGTCSLH
TTASTLDDDGNYTIMAANPQGRVSCTGRLMVQAVNQRGRSPRSP---------------------
PIQERFFRPHFLQAPGDLTVQEGKLCRMDCKVSGLPTPDLSWQLDGKPIRPDSAHKMLVRENGVHSLI
IEPVTSRDAGIYTCIATNRAGQNSFNLELVVAAKEAHK

- Ig3-4Δ6

SNANATAPFFEMKLKHYKIFEGMPVTFTCRVAGNPKPKIYWFKDGKQISPKSDHYTIQRDLDGTCSLH
TTASTLDDDGNYTIMAANPQGRVSCTGRLMVQAVNQRGRSPRSPSGHPHARRPRSRSRD------
PIQERFFRPHFLQAPGDLTVQEGKLCRMDCKVSGLPTPDLSWQLDGKPIRPDSAHKMLVRENGVHSLI
IEPVTSRDAGIYTCIATNRAGQNSFNLELVVAAKEAHK

- Ig3-4Δ4-SGHP

SNANATAPFFEMKLKHYKIFEGMPVTFTCRVAGNPKPKIYWFKDGKQISPKSDHYTIQRDLDGTCSLH
TTASTLDDDGNYTIMAANPQGRVSCTGRLMVQAVNQRGRSPRSP----
HARRPRSRSRDSGDENEPIQERFFRPHFLQAPGDLTVQEGKLCRMDCKVSGLPTPDLSWQLDGKPIRP
DSAHKMLVRENGVHSLIIEPVTSRDAGIYTCIATNRAGQNSFNLELVVAAKEAHK

- Ig3-4ΔHPHA

SNANATAPFFEMKLKHYKIFEGMPVTFTCRVAGNPKPKIYWFKDGKQISPKSDHYTIQRDLDGTCSLH
TTASTLDDDGNYTIMAANPQGRVSCTGRLMVQAVNQRGRSPRSPSG----

RRPRSRSRDSGDENEPIQERFFRPHFLQAPGDLTVQEGKLCRMDCKVSGLPTPDLSWQLDGKPIRPDS
AHKMLVRENGVHSLIIEPVTSRDAGIYTCIATNRAGQNSFNLELVVAAKEAHK

- Ig3-4ΔPHAR

SNANATAPFFEMKLKHYKIFEGMPVTFTCRVAGNPKPKIYWFKDGKQISPKSDHYTIQRDLDGTCSLH
TTASTLDDDGNYTIMAANPQGRVSCTGRLMVQAVNQRGRSPRSPSGH----
RPRSRSRDSGDENEPIQERFFRPHFLQAPGDLTVQEGKLCRMDCKVSGLPTPDLSWQLDGKPIRPDSA
HKMLVRENGVHSLIIEPVTSRDAGIYTCIATNRAGQNSFNLELVVAAKEAHK

- Ig3-34L-Ig5

SNANATAPFFEMKLKHYKIFEGMPVTFTCRVAGNPKPKIYWFKDGKQISPKSDHYTIQRDLDGTCSLH
TTASTLDDDGNYTIMAANPQGRVSCTGRLMVQAVNQRGRSPRSPSGHPHARRPRSRSRDSGDENEPIQ
ERFFRPVFMEKLQNTGVADGYPVRLECRVSGVPPPQIFWKKENESLTHSTERVSMHQDNHGYICLLIQ
GATKEDAGWYTVSAKNEAGIVSCTARLDVY

- Ig3-45L-Ig5

SNANATAPFFEMKLKHYKIFEGMPVTFTCRVAGNPKPKIYWFKDGKQISPKSDHYTIQRDLDGTCSLH
TTASTLDDDGNYTIMAANPQGRVSCTGRLMVQAKEAHKAPVFMEKLQNTGVADGYPVRLECRVSGVPP
PQIFWKKENESLTHSTERVSMHQDNHGYICLLIQGATKEDAGWYTVSAKNEAGIVSCTARLDVY

- Ig3-45L-Ig4

SNANATAPFFEMKLKHYKIFEGMPVTFTCRVAGNPKPKIYWFKDGKQISPKSDHYTIQRDLDGTCSLH
TTASTLDDDGNYTIMAANPQGRVSCTGRLMVQ**AKEAHKA**PHFLQAPGDLTVQEGKLCRMDCKVSGLPT
PDLSWQLDGKPIRPDSAHKMLVRENGVHSLIIEPVTSRDAGIYTCIATNRAGQNSFNLELVVAAKEAH
K

- Ig3-4 LinkerScramble

SNANATAPFFEMKLKHYKIFEGMPVTFTCRVAGNPKPKIYWFKDGKQISPKSDHYTIQRDLDGTCSLH
TTASTLDDDGNYTIMAANPQGRVSCTGRLMVQ**DNHERERRIRERSRFDPSAPPRRASQGSRSHFGRSN**
**VGQPP**PHFLQAPGDLTVQEGKLCRMDCKVSGLPTPDLSWQLDGKPIRPDSAHKMLVRENGVHSLIIEP
VTSRDAGIYTCIATNRAGQNSFNLELVVAAKEAHK

- Ig3-4 RLinkerA

SNANATAPFFEMKLKHYKIFEGMPVTFTCRVAGNPKPKIYWFKDGKQISPKSDHYTIQRDLDGTCSLH
TTASTLDDDGNYTIMAANPQGRVSCTGRLMVQAVNQ**A**G**A**SP**A**SPSGHPHA**AA**PAS**A**S**A**DSGDENEPIQE
**A**FF**A**PHFLQAPGDLTVQEGKLCRMDCKVSGLPTPDLSWQLDGKPIRPDSAHKMLVRENGVHSLIIEPV
TSRDAGIYTCIATNRAGQNSFNLELVVAAKEAHK

- Ig3-4 DELinkerA

SNANATAPFFEMKLKHYKIFEGMPVTFTCRVAGNPKPKIYWFKDGKQISPKSDHYTIQRDLDGTCSLH
TTASTLDDDGNYTIMAANPQGRVSCTGRLMVQAVNQRGRSPRSPSGHPHARRPRSRSR**A**SG**AA**NEPIQ**A**
RFFRPHFLQAPGDLTVQEGKLCRMDCKVSGLPTPDLSWQLDGKPIRPDSAHKMLVRENGVHSLIIEPV
TSRDAGIYTCIATNRAGQNSFNLELVVAAKEAHK

**Supplemental Note 2:** Primer sequences

- Ig3-4Δ4-SGHP—one primer only
  - GCAGTCCCCGCTCTCCCCATGCCAGAAGGCCTCGCTC
- Ig3-4ΔHPHA
  - Forward: TCCCCGCTCTCCCTCAGGCAGAAGGCCTCGCTCTCG
  - Reverse: CCGTGATCGAGAGCGAGGCCTTCTGCCTGAGGGAGAGCGG
- Ig3-4ΔPHAR
  - Forward: GCTCTCCCTCAGGCCATAGGCCTCGCTCTCGATCA
  - Reverse: TGATCGAGAGCGAGGCCTATGGCCTGAGGGAGAGC
- Ig3-4Δ21—one primer only
  - CAGTCCCCGCTCTCCCCCCATTCAGGAGCGATTCTTCAG
- Ig3-4Δ6—two primers
  - Forward: CGCTCTCGATCACGGGACCCCATTCAGGAGCGATTCTTCA
  - Reverse: TGAAGAATCGCTCCTGAATGGGGTCCCGTGATCGAGAGCG
- Ig3-4-RLinkerA
  - Forward:
    CTGCATcagcggacagtggagatgaaaacgagcccattcaggaggcattcttcgcaCCT
    CACTTCCTGCAGGCT
  - Reverse:
    GTACAGGCTGTCAACCAAgcaggcgccagtcccgcctctccctcaggccatcctcatgc
    cgcagcgccTGCCT
- Ig3-4-DELinkerA
  - Forward : acgcgcccattcaggcgCGATTCTTCAGACCTC
  - Reverse: ttgcagctccactggcCCGTGATCGAGAGCGAG
- Ig3-4-LinkerScramble
  - Forward:
    TCGACGGGCCTCacaaggctctcgatcacatttcggcagaagtaacgtcggacagcctc
    ccCCTCACTTCCTGCAGGCT
  - Reverse:
    CAGGAAGGCTAATGGTACAGgacaaccatgaaagagagcgccgcattagagagaggagt
    cgcttcgatcCCCCCTCGTCTCC
- Ig3-45L-Ig4
  - Forward: cacaaggccCCTCACTTCCTGCAGGCT
  - Reverse: tgcttccttAGCCCTGTACCATTAGCCTTC
- Ig3-45L-Ig5
  - Forward: GCTAAGGAAGCACACAAG
  - Reverse: CTGTACCATTAGCCTTCC
- Ig3-34L-Ig5
  - Forward: gaacacgggggttgctgatGGATACCCAGTGCGGCTA
  - Reverse: tgtagcttctccataaacacAGGTCTGAAGAATCGCTCC

Supplemental Materials

**Supplementary Note 3. SAXS data deposition link**

Link to SASBDB (Small Angle Scattering Biological Data Bank)

ID: DX77 (Ig3-45L-Ig4) along with previously published DVE6 (Ig3), DVD6 (Ig4), and DVF6 (Ig3-4 WT).
